# Widely Used Calculations of Intracranial Volume Provide Inaccurate Estimates in Developmental Cohorts: Evidence and Recommendations for the Neuroimaging Community

**DOI:** 10.64898/2026.10.03.751882

**Authors:** Amalia M. Skyberg, Ava J. Reck, Jolinda Smith, Benjamin T. Newman, Madeleine N. Goldberg, T. Jason Druzgal, John Van Horn, Jennifer H. Pfeifer

## Abstract

Intracranial volume (ICV) is estimated as part of automated neuroimaging processing pipelines and commonly used to account for and characterize cranial size. FreeSurfer’s estimated intracranial volume (eTIV) produces unreliable estimates in developmental samples when using versions prior to FreeSurfer 8, which longitudinal cohorts reveal to be biologically implausible. Across two different adolescent datasets, we document eTIV changes of up to 39% between 1.5 - 2 year sessions for FreeSurfer 7, with errors more pronounced on Siemens scanners and in female participants. The errors are derived from failures of an early registration step, which was upgraded to a deep learning registration algorithm in FreeSurfer 8, where these errors are markedly reduced and produce eTIV estimates comparable to an independent estimation method, AssemblyNet. However, the widespread use of FreeSurfer 7 raises concern about developmental neuroimaging studies that utilize eTIV without quality control. Recommendations for future research and considerations for past research in the field are discussed.

---

Intracranial volume (ICV) refers to the total volume contained within the cranial vault, including brain tissue, cerebrospinal fluid, meninges, and the internal spaces bounded by the skull. In neuroimaging studies, ICV adjustment is commonly used to account for normal variation in head size, since larger people, on average, tend to have larger cranial vaults and larger absolute brain structures. This is important to ensure that volumetric differences reflect meaningful regional anatomy rather than premorbid cranial size. Once computed, ICV can be used as a dependent variable or included as a covariate used to normalize regional volumes, incorporated through residualization or allometric scaling to distinguish regional effects from global differences in cranial capacity. The widespread utility of ICV makes the reliability of its estimation an essential concern.

Due to the insufficient signal to visualize cortical bone in T1-weighted images, ICV cannot be measured by delineating the inner table of the skull, and must be estimated. Most estimation methods fall into two categories: segmentation-based methods, which estimate ICV by summing segmented gray matter, white matter, and cerebrospinal fluid volumes (used by SPM (Friston et al., 2007)) and registration-based, which infer ICV from the transformation required to align an individual’s brain to a template (used by FSL (Jenkinson et al., 2012) and Freesurfer (Fischl, 2012)).

In developmental cognitive neuroscience, FreeSurfer is one of the most commonly used tools for analyzing and visualizing structural and functional neuroimaging data across both cross-sectional and longitudinal designs. FreeSurfer’s ICV estimate is widely used in the published literature. For example, a Google Scholar search for “intracranial volume” AND “freesurfer” returns 16,300 articles (6,410 if restricted to include the terms “adolescent” OR “children”). Moreover, FreeSurfer is the standard analysis pipeline for structural data in landmark population datasets such as Adolescent Brain and Cognitive Development (ABCD) Study® (Casey et al., 2018), UK BioBank (Miller et al., 2016), and Human Connectome Projects (HCP; (Glasser et al., 2013)).

FreeSurfer versions 4-7 report an estimated ICV (eTIV) using an atlas-based normalization procedure, derived from the automated affine registration of the anatomic image to the MNI305 atlas in Talaraich space (Buckner et al., 2004). This method works under the assumption that the scaling factor needed to register the participant’s brain to a template is identical to the scaling factor between the skull and the template skull. Under this assumption, the reciprocal of the determinant of the transformation matrix will be proportional to the ICV (specifically, eTIV = 1755cm^3^/ASF). While there are other methods for obtaining ICV, including segmentation-based (Puonti et al., 2016) and machine learning methods (Coupé et al., 2020), eTIV is one of the most accessible and widely used metrics. eTIV is automatically produced by FreeSurfer’s recon-all pipeline and is distributed in released tabulated datasets for landmark population datasets as noted above. It should be noted that the Freesurfer version changes with different ABCD releases, such as Freesurfer 5.4 for ABCD release 2.0.1, and Freesurfer 7.1.1 for ABCD release 4.0)

The current study was motivated by unexpected, biologically implausible longitudinal patterns observed when using eTIV derived from FreeSurfer 7.3, included as part of the fMRIPrep pipeline (Esteban et al., 2019). In a dataset of 217 adolescent girls assessed up to six times over eight years (Barendse et al., 2020), eTIV produced anomalous structural trajectories when used as a dependent variable, and significantly altered developmental trajectories when used as a covariate. Specifically, we observed that adding cross-sectionally derived eTIV as a covariate into a mixed effect model using age to predict hippocampal volume derived from fMRIPrep’s default FreeSurfer pipeline transformed a significant linear trajectory to a significant cubic trajectory. This prompted us to look at eTIV by age, which led to the discovery of the eTIV errors (see Figure 1). Irregularities were similarly observed in analyses of eTIV from the ABCD Study across two waves.

**Figure 1.**
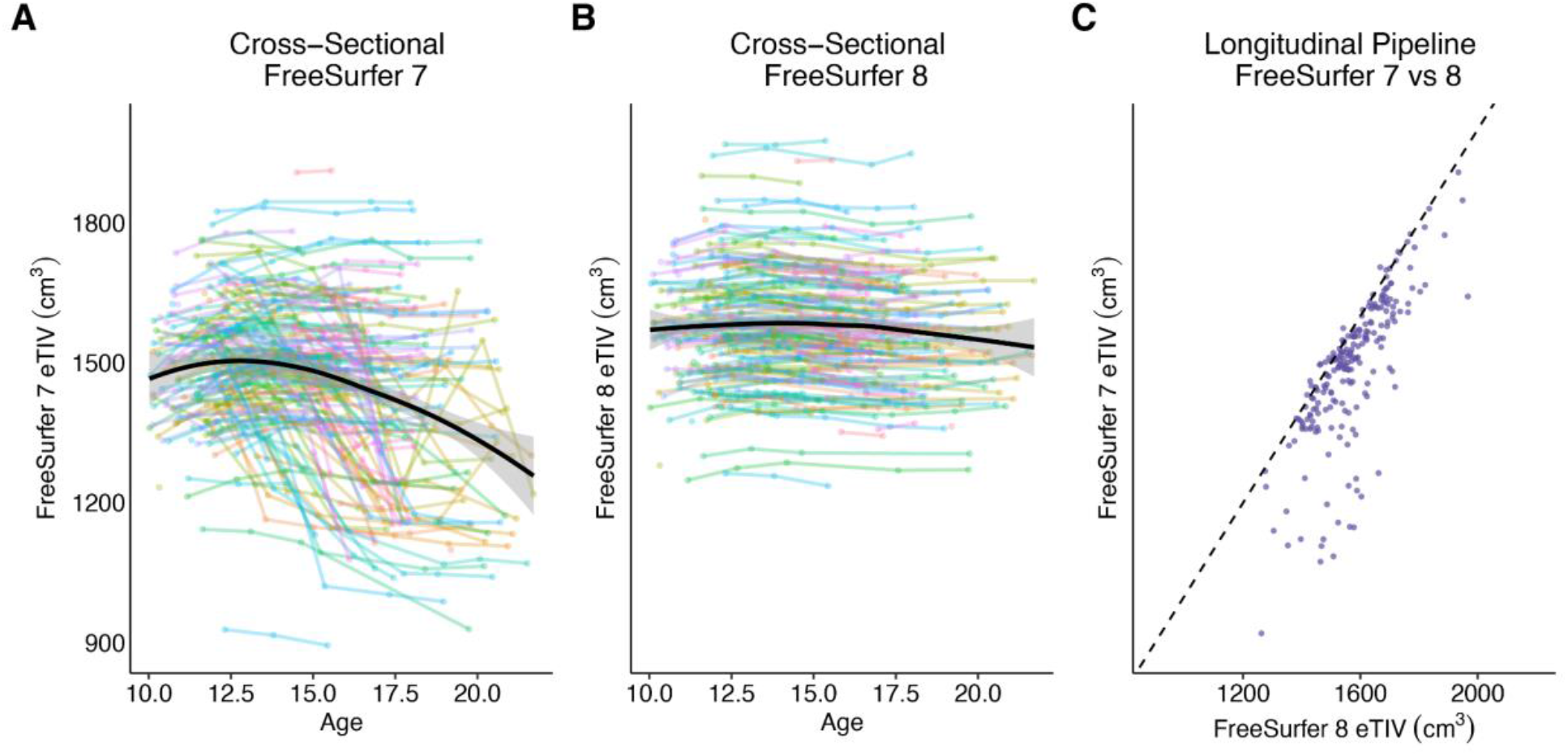
Developmental trajectories of eTIV in the TAG cohort derived from FreeSurfer 7 and FreeSurfer 8. In panels A and B, each colored line represents the eTIV value of an individual over time, highlighting the implausible trajectories produced using FreeSurfer 7. The solid black line with the shaded 95% confidence interval indicates the overall trend in the cohort. Panel C shows the correspondence between the single eTIV estimate produced using the longitudinal version of the pipelines. The black dashed line represents the expected relationship under perfect proportional agreement (slope = 1).

These observations prompted concerns about the reliability of eTIV estimates in longitudinal samples and their potential to introduce systematic bias into morphometric analyses. While minor variation in eTIV has been described in lifespan and clinical aging literatures (Klasson et al., 2018; Koike et al., 2025; Nerland et al., 2022), this issue remains largely unrecognized in other subfields of cognitive neuroscience. Furthermore, the degree of biologically implausible variation in eTIV we identify in developmental samples is far outside the bounds described in these papers. Continued reliance on eTIV thus potentially compromises the validity of a substantial portion of the existing literature to an unknown degree. Therefore, the purpose of this paper is to document and characterize erroneous longitudinal behavior of FreeSurfer-derived eTIV across two developmental datasets, as well as describe recommendations to eliminate eTIV as a source of error in cognitive neuroscience research.

## Methods

### Data used in the discovery of biologically implausible eTIV trajectories

Data were first obtained from the Transitions in Adolescent Girls (TAG) study (Barendse et al., 2020). The sample comprised two recruitment bursts: an original cohort of 172 participants (mean age = 11.54 years at study intake; range = 10.00-12.99, SD = 0.81) and a replenishment cohort of 45 participants (mean age = 15.32 years at 14.03-17.00; range, 0.80), yielding a total sample of 217 youth, all assigned female at birth. Participants were scanned up to six times (mean intersession interval = 1.7 years) on a Siemens Skyra scanner. T1w MPRAGE scans (voxel size 1x1x1mm^3^, FOV: 256x256x256mm) were preprocessed using fMRIPrep 25.2.5, which includes FreeSurfer 7.3.2 in its pipeline. The images were also preprocessed with FreeSurfer 8.2.0 (released in March 2026) and AssemblyNet (Coupe et al., 2020), a neural network-based segmentation pipeline. As FreeSurfer’s longitudinal pipeline estimates a single ICV, the ICV measures for the current study were extracted from each scan after being processed through the cross-sectional pipeline, in addition to being processed through the longitudinal pipeline. The estimated ICVs were compared for the three pipelines (FreeSurfer V7, FreeSurfer V8, and AssemblyNet). In addition, we reran FreeSurfer 7 using a corrected transformation and included the --no--randomness flag to enable strict comparison within version 7 in this dataset. All study procedures were approved by the University Institutional Review Board and were conducted in accordance with the ethical principles of the Declaration of Helsinki. Written informed consent was obtained from all participants or their legal guardians, and assent was obtained from minors prior to participation.

### Secondary data used to validate the observed eTIV errors from FreeSurfer 7

Data were also obtained from the ABCD 4.0 minimally processed release (the data from the ABCD Study release 4.0 utilized FreeSurfer 7.1.1). While an older release, more recent releases were not available during the data processing period of this paper due to access restrictions. Informed consent/assent was obtained, and all study procedures were approved by a central Institutional Review Board and were conducted in accordance with the ethical principles of the Declaration of Helsinki. Twenty percent of subjects were randomly selected from each site for further analysis to maintain demographic and population distributions. This subsample totaled 2,529 individuals (1,213 female), with a mean age of 9.90 years at study intake (range = 8.92-11.00, SD = 0.63) who were scanned one or two times (mean intersession interval = 2.0 years). T1w MPRAGE images (voxel size 1x1x1mm^3^, FOV: 256x256x256mm) were downloaded from the NIMH data repository (NDA) and were not preprocessed, defaced, or anonymized in any way prior to analysis. All images were analyzed through the recon-all command in FreeSurfer 8.1.0 and separately with AssemblyNet 1.0.0 both implemented in singularity containers on a linux HPC environment. Additionally, we investigated whether scanner type or sex accounted for changes between waves in ABCD. Lastly, because studies have shown that scanner-induced variance is present across multiple types of neuroimaging data and can exceed variability attributable to biological factors such as age and sex, longitudinal harmonization using ComBat was employed on the ABCD data (Beer et al., 2020; Dudley et al., 2023).

## Results

### Comparison of within individual changes in ICV utilizing different pipelines

In both datasets, FreeSurfer 8 and AssemblyNet demonstrated stable ICV estimates between scan sessions with mean absolute percentage changes hovering around 1%, whereas FreeSurfer 7 showed approximately three times greater mean absolute percentage change, summarized in Table 1. Briefly, in the TAG sample, FreeSurfer 7 eTIV values varied by up to 521 cm^3^ between sessions (mean change = 51 cm^3^). In contrast, FreeSurfer 8 showed a maximum session-to-session eTIV difference of 63 cm^3^ (mean change < 14 cm^3^). Similarly, in the ABCD sample, FreeSurfer 7 eTIV values varied by up to 368 cm^3^ between sessions (mean change = 37 cm^3^). FreeSurfer 8 showed a maximum session-to-session eTIV difference of 74 cm^3^ (mean change 21 cm^3^). Comparison of the single estimate produced from the longitudinal version of the pipeline in FreeSurfer 7 and FreeSurfer 8 also demonstrated poor correspondence (see Figure 1) indicating that the error is pervasive in either approach.

**Table 1.**
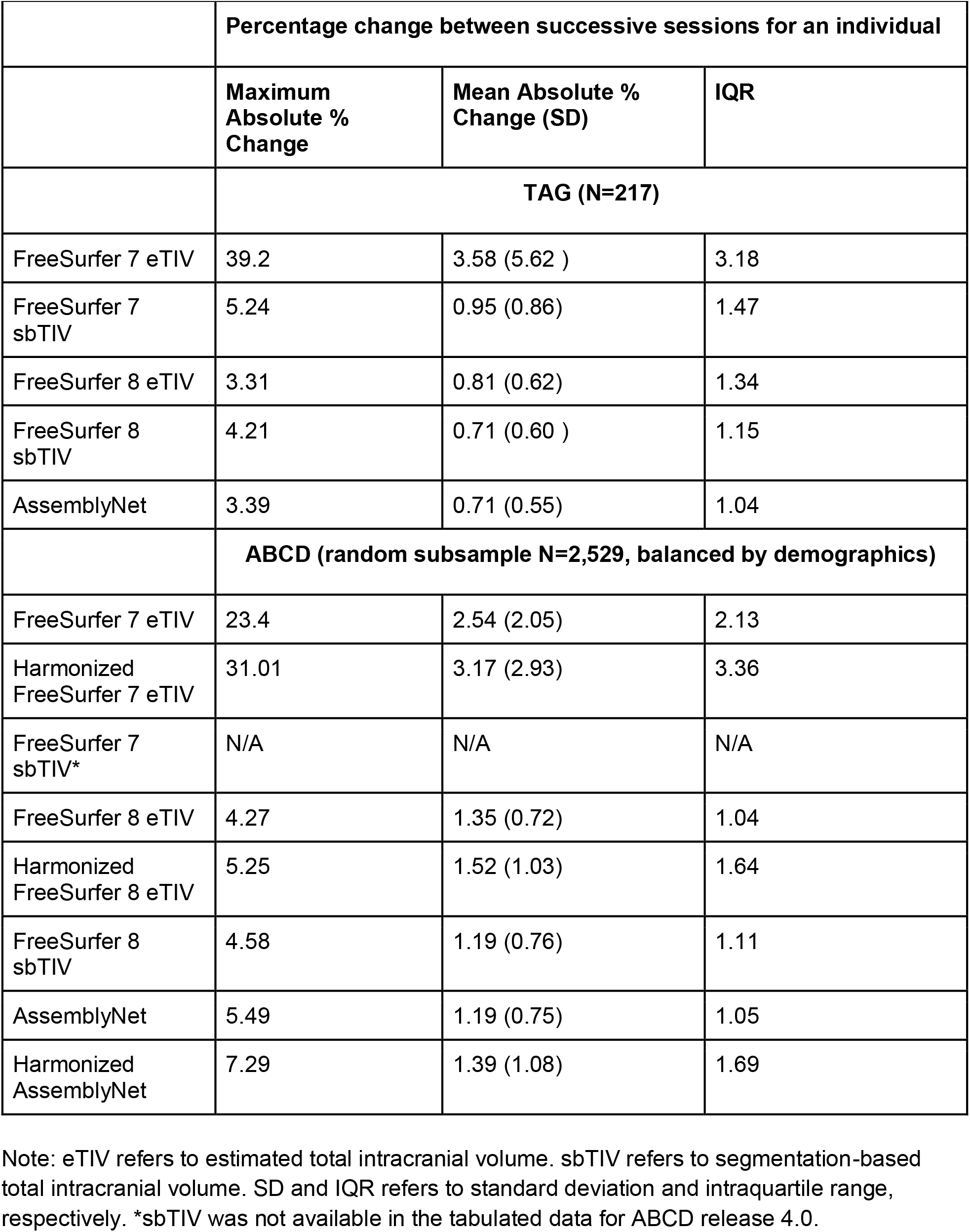
Percentage change metrics between successive sessions in TAG and ABCD samples.

|  | **Percentage change between successive sessions for an individual** |  |  |
| --- | --- | --- | --- |
|  | **Maximum Absolute % Change** | **Mean Absolute % Change (SD)** | **IQR** |
|  | **TAG (N=217)** |  |  |
| FreeSurfer 7 eTIV | 39.2 | 3.58 (5.62 ) | 3.18 |
| FreeSurfer 7 sbTIV | 5.24 | 0.95 (0.86) | 1.47 |
| FreeSurfer 8 eTIV | 3.31 | 0.81 (0.62) | 1.34 |
| FreeSurfer 8 sbTIV | 4.21 | 0.71 (0.60 ) | 1.15 |
| AssemblyNet | 3.39 | 0.71 (0.55) | 1.04 |
|  | **ABCD (random subsample N=2,529, balanced by demographics)** |  |  |
| FreeSurfer 7 eTIV | 23.4 | 2.54 (2.05) | 2.13 |
| Harmonized FreeSurfer 7 eTIV | 31.01 | 3.17 (2.93) | 3.36 |
| FreeSurfer 7 sbTIV* | N/A | N/A | N/A |
| FreeSurfer 8 eTIV | 4.27 | 1.35 (0.72) | 1.04 |
| Harmonized FreeSurfer 8 eTIV | 5.25 | 1.52 (1.03) | 1.64 |
| FreeSurfer 8 sbTIV | 4.58 | 1.19 (0.76) | 1.11 |
| AssemblyNet | 5.49 | 1.19 (0.75) | 1.05 |
| Harmonized AssemblyNet | 7.29 | 1.39 (1.08) | 1.69 |

Note: eTIV refers to estimated total intracranial volume. sbTIV refers to segmentation-based total intracranial volume. SD and IQR refers to standard deviation and intraquartile range, respectively. \*sbTIV was not available in the tabulated data for ABCD release 4.0.

Review of FreeSurfer documentation and visual inspection of extreme outliers in terms of percent change in session-to-session eTIV for both TAG and ABCD revealed that the error in FreeSurfer 7 originates from the registration step early in the pipeline. As shown in Figure 2, the downstream consequences of poor registration can be visualized in the brain image (brain.mgz) and the cortical surface outline (lh.ribbon.mgz and rh.ribbon.mgz) outputs. FreeSurfer 8 documentation indicates that the registration tool used in FreeSurfer 7 was changed in FreeSurfer 8 to SynthMorph (Hoffmann et al., 2022), a more robust and efficient deep learning algorithm. Correspondence of ICV estimates across pipelines (FreeSurfer 7, FreeSurfer 8, and AssemblyNet) are presented in Figure 3.

**Figure 2.**
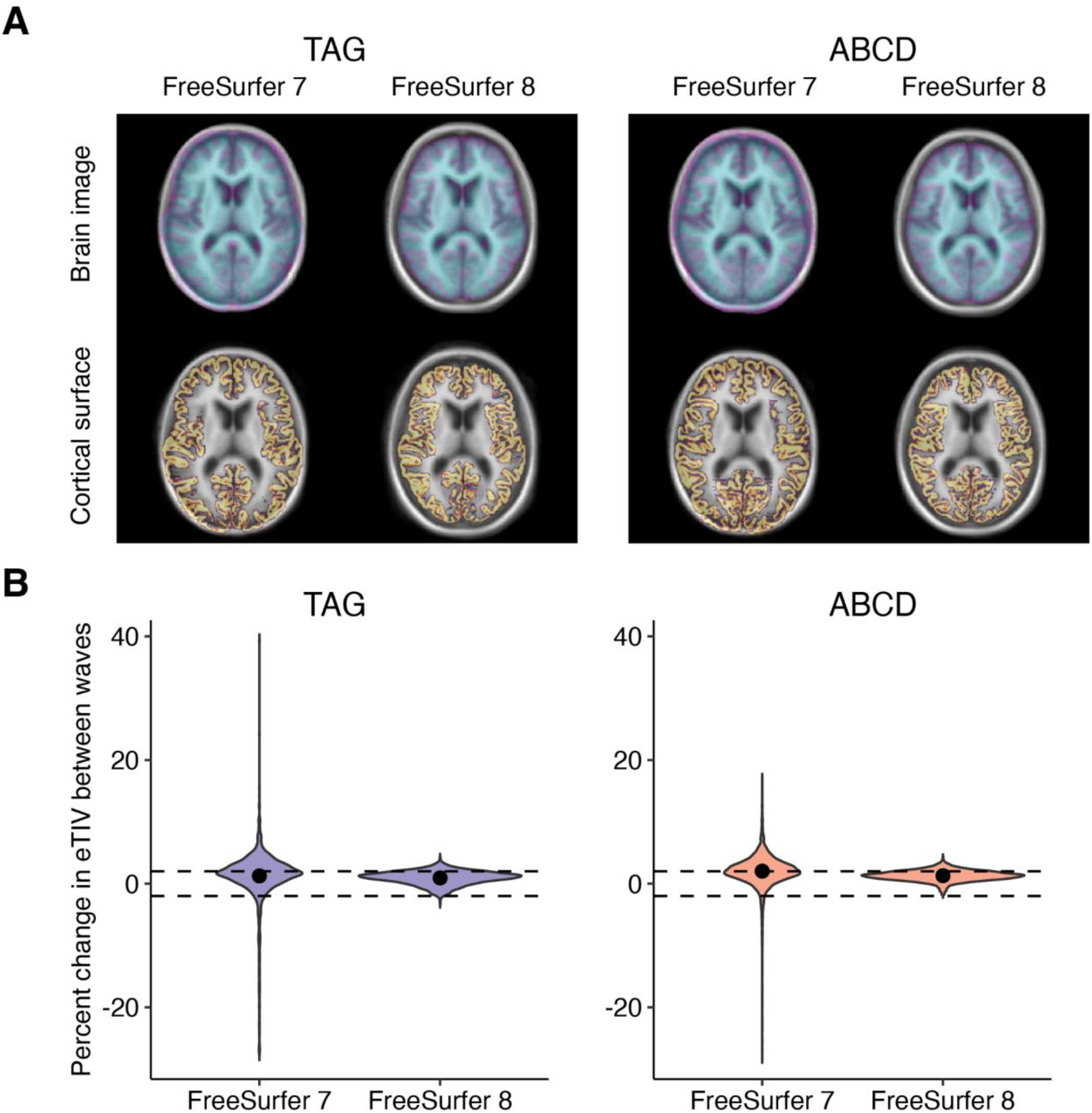
Panel A visualizes failed registration in a single subject from each of TAG and ABCD, overlaid on the fsaverage/orig.mgz. Top row depicts the brain image (brain.mgz) transformed to MNI space (mni305.cor.mgz) using talairach.xfm; bottom row depicts the cortical surface outline (lh.ribbon.mgz and rh.ribbon.mgz) transformed to MNI space (mni305.cor.mgz) using talairach.xfm. Panel B depicts violin plots of percent change in eTIV between each wave derived from FreeSurfer 7 and FreeSurfer 8. The filled circle represents the mean. The dashed lines at ± 2% represent generous boundaries in plausible eTIV percent change due to developmental change over an 18-24 month period (Caspi et al., 2020) or measurement error.

**Figure 3.**
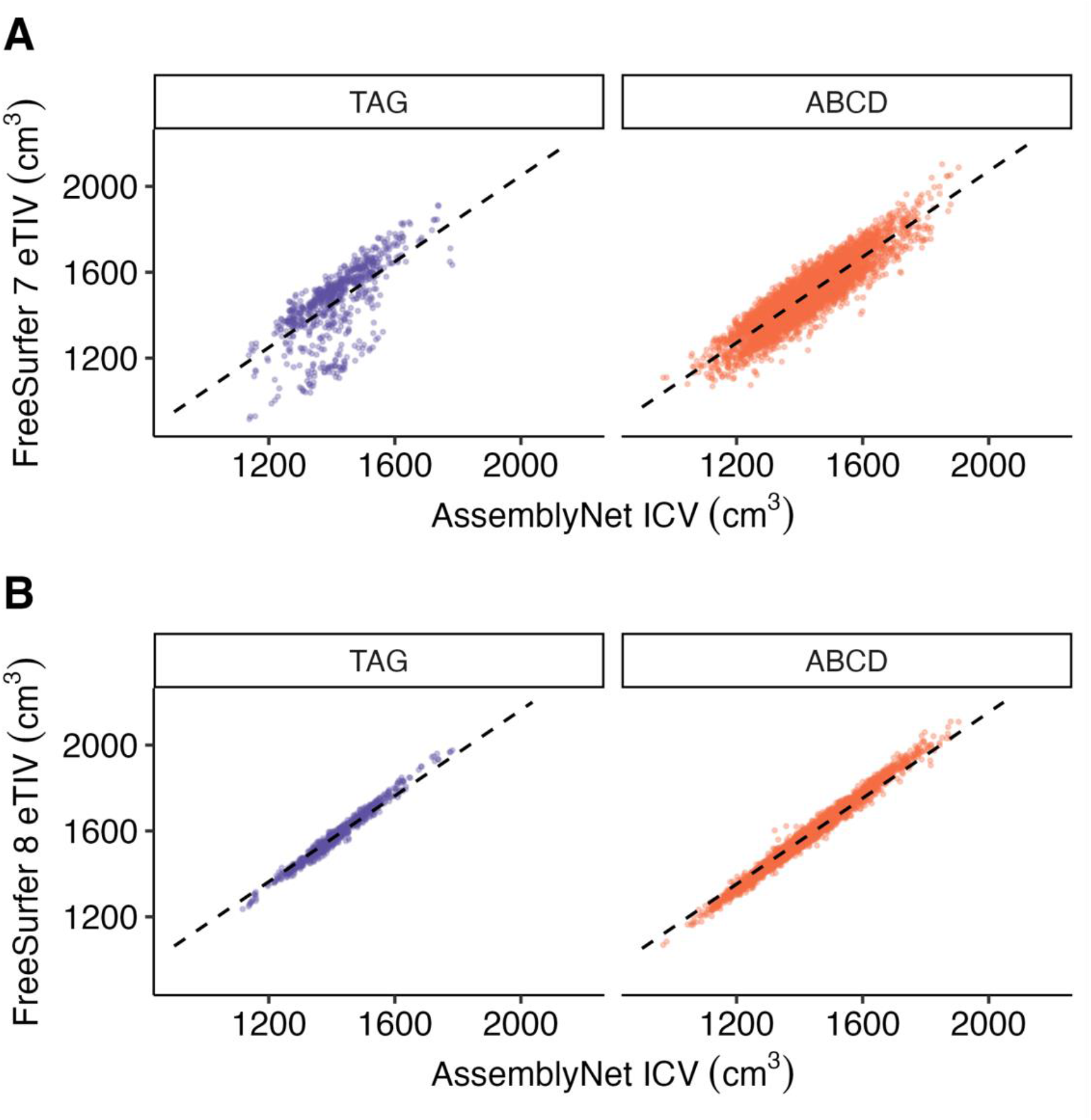
Correspondence between ICV across analysis pipelines. Panels A and B depict correspondence between ICV derived from FreeSurfer versions 7 and 8, respectively, and AssemblyNet across TAG and ABCD samples. The black dashed lines represent the expected relationship under perfect proportional agreement (slope = 1).

### Reliability within and across FreeSurfer versions

eTIV is a global brain metric, which leaves unaddressed the question of whether the unreliability of the Talairach transformation early in FreeSurfer 7 affects the regional volume, surface area, or cortical thickness outputs of recon-all. Supplementary Figures X/Y/Z visualize the reliability of rank ordering outputs between FreeSurfer 7 with and without correcting the transformation while holding the random seed constant, as well as between FreeSurfer 7 (corrected) and FreeSurfer 8. Correspondence between FreeSurfer versions is not expected to be perfect due to continual processing improvements (Haddad et al., 2022). In general, the reliability between versions 7 and 8 is at least as good as that previously demonstrated between versions 6 and 7, if not better across many regions that were previously unreliable. However, regions that are highly susceptible to artifacts figure prominently in the category of regions that demonstrate lower reliability across versions, or when accounting for poor Talairach transformations. On the other hand, eTIV reliability seems to be more clearly due to the transformation, as using a corrected transform increased the reliability from .72 to .99; it is the only metric where there is a bigger difference in reliability between using the corrected and uncorrected transforms, than between changing versions of FreeSurfer.

### Biological sex and scanner manufacturer contribute to the severity of eTIV changes

Figures 2 and 3 indicate that the TAG study had more extreme changes in FreeSurfer 7 eTIV between waves than was observed in ABCD. We investigated whether scanner type or sex could account for changes between waves in the full ABCD dataset. A one-way ANOVA revealed a significant effect of scanner manufacturer on percent change in FreeSurfer 7 eTIV, F(2, 15549) = 115.60, *p* < 0.001. A post hoc Tukey test indicated that Siemens scanners exhibited more negative changes between the two waves compared to both GE and Phillips scanners (mean differences = -0.61% and -0.65%, ps < 0.001, respectively; Figure 4).

**Figure 4.**
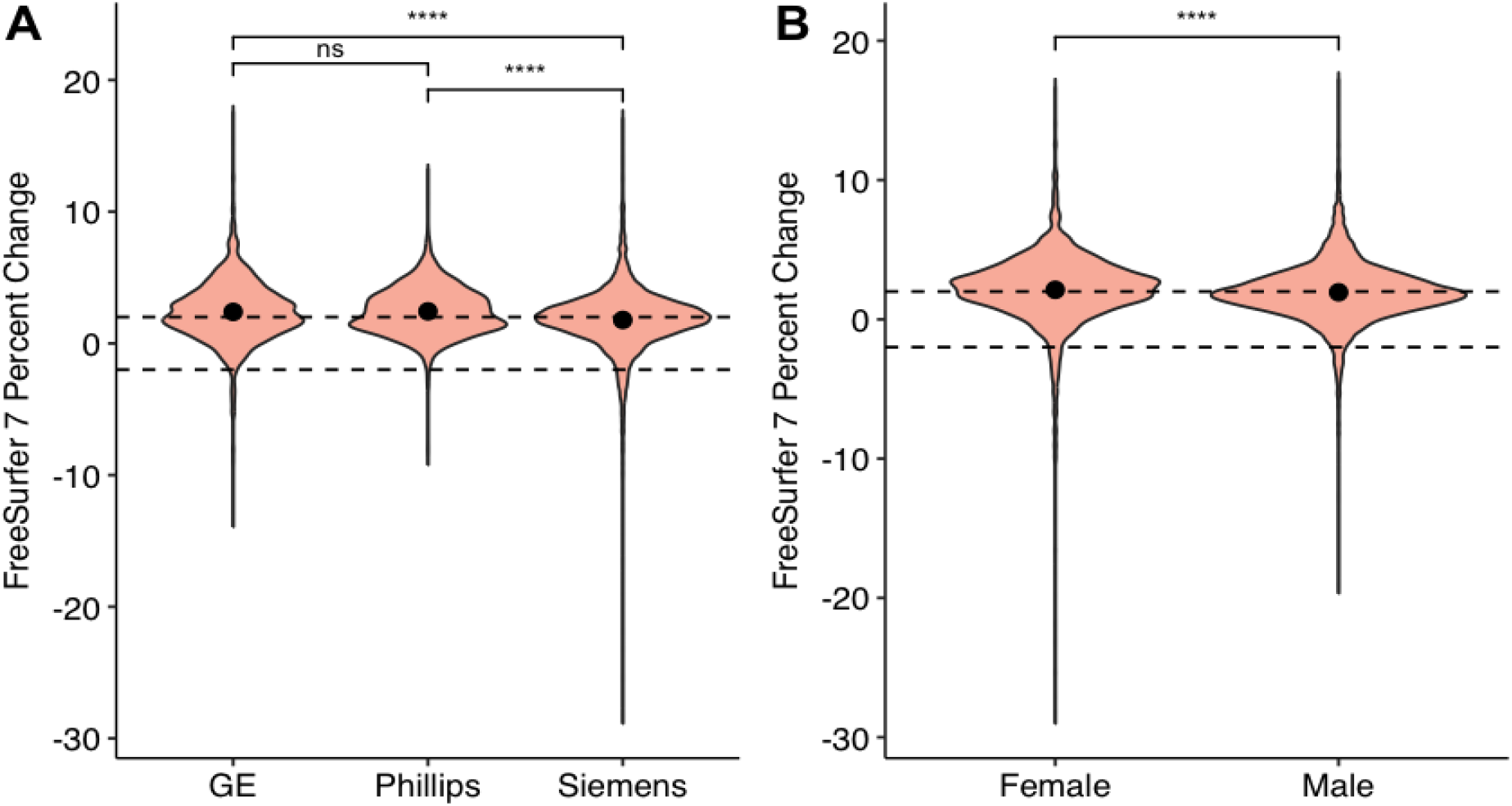
Violin plot visualization of the results from **A)** the one-way ANOVA for scanner manufacturer and **B)** the t-test for sex differences in observed percent changes in FreeSurfer 7 eTIV estimates. Ns = non significant, **** indicates *p* < 0.001.

Additionally, female participants exhibited significantly more changes between the two waves (t(13,320) = 4.32, *p* < 0.001; Figure 4). We suspected that this was more common in female participants because of smaller brain size, and indeed when controlling for ICV (using AssemblyNet estimates), the association between sex and percent change in eTIV between waves was no longer significant (F(1, 3395) = 2.65, *p* = 0.104) but ICV was (F(1, 3395) = 6.28, *p* = 0.012). This likely helps to explain the substantial degree of eTIV errors in the TAG study as all participants were assigned female at birth and exclusively scanned on a Siemens scanner.

### Scanner software updates are not associated with within-person eTIV estimate variability

To check if scanner software updates over the study time course contributed to within-person variability in eTIV estimates, model comparison including software version was tested in the TAG cohort. Scanner software version did not explain within-person variability (B = -0.83 cm^3^, *p* > 0.05) and including scanner software version did not improve model fit beyond FreeSurfer version alone (F(1,363) = 0.02, *p* > 0.05). These results suggest that eTIV estimate instability between scan sessions is attributable to FreeSurfer processing pipeline version rather than MRI scanner software upgrades.

### Harmonizing across sites does not improve within-person eTIV estimate reliability

Because significant differences were observed by scanner manufacturers, longCombat harmonization was employed on the ABCD subset to query if harmonization across sites resolved the observed within-person changes in eTIV estimates. As seen in Table 1, results of absolute change from the harmonization approach were on the same order of magnitude for FreeSurfer 7, FreeSurfer 8, and AssemblyNet. These results further highlight an error in the pipeline, and while harmonization is intended to address systematic technical differences between scanners, it does not necessarily correct errors arising from the estimation of a measure.

## Discussion

The present study demonstrates that FreeSurfer-derived eTIV, from versions prior to FreeSurfer 8, can produce unreliable ICV estimates in developmental samples over an 18-24 month period. Anomalous ICVs were pronounced in FreeSurfer 7, markedly reduced in FreeSurfer 8, and largely absent when using an independent segmentation-based estimate of intracranial volume (AssemblyNet). In addition, eTIV errors were significantly more common on Siemens scanners and in female youth, and not attributable to software upgrades.Despite recent work emphasizing the need for careful quality control in harmonizing multi-site ABCD data (Kraft et al., 2024), our findings further suggest that harmonization does not resolve within-person eTIV instability when the source of error lies in the underlying volume estimation rather than systematic scanner differences.

### Considerations for recon-all output from earlier FreeSurfer versions

Several cautions follow from the current results. First, prior developmental studies that relied on FreeSurfer-derived eTIV prior to version 8 without explicit quality control of the registration step should be interpreted with caution. Because this processing step occurs early in the pipeline and is not routinely inspected in most workflows, undetected registration failures likely propagated into estimates of ICV. Consequently, some reported developmental changes in ICV (e.g., steep nonlinear increases from birth to age six (Alex et al., 2024); apparent stabilization in late adolescence (Mills et al., 2016)) may partly reflect measurement instability rather than true neurodevelopmental change.

Second, the discovery of this error through the inclusion of eTIV as a covariate, which made linear volumetric trajectories significantly nonlinear, also has important implications for developmental neuroimaging, where ICV is routinely modeled as a nuisance variable to account for global brain size. This might explain the contradicting effects of ICV vs WBV correction in the literature (Mills et al., 2016) as well as the more recent demonstrations that proportional ICV correction can affect predictive accuracies (Dhamala et al., 2022). These data show that not only are the eTIV estimates noisy, but they are subject to systematic bias which is not improved by harmonization approaches. Left unchecked, this has the potential to distort growth models, sex comparisons, and brain-behavior associations.

The findings described in the current paper raise substantial concern regarding the use of eTIV in neuroimaging studies. Methodologically, eTIV depends on successful affine registration of the T1-weighted image to a standard template, a step that occurs early in the FreeSurfer pipeline and may not be routinely inspected or reported. Sagely, in the context of discussing an error in automated eTIV over two decades ago, Buckner and colleagues (2004) reminded readers that “automated procedures need not, and should not, be unsupervised.” Because eTIV is derived directly from the transformation matrix, it is uniquely sensitive to errors in the scaling factor of this transform. Nevertheless, because this calculation occurs early in the recon-all pipeline, errors in the talairach.xfm matrix could potentially influence subsequent processing steps and, in-turn, affect other FreeSurfer derived measures. Our analyses suggest that some subcortical and cortical recon-all outputs, previously implicated in susceptibility and reliability concerns, may be moderately affected (Supplementary Figure 1A and 1B).

### Reinforcing data and analysis quality in an era of big data and automated pipelines

These findings highlight persistent challenges for quality control in large datasets. While large consortia datasets have transformed the field by enabling unprecedented statistical power and accessibility, increasing sample sizes make routine visual quality control impractical. Additionally, greater accessibility of these large datasets means that researchers frequently engage more with tabulated derivatives rather than the raw imaging data. Large-scale archival datasets such as ABCD and HCP-D provide eTIV from previous FreeSurfer versions in tabulated datasets without visual examination. Until these consortia update their pipeline and data releases, researchers should not use previously released eTIV data without first checking for unreliable estimates. Likewise, analysis pipelines that utilize eTIV (e.g., ENIGMA BrainAge; Han et al., 2022) need to be revisited. Additionally, we show that some regions continue to be more susceptible to changes between FreeSurfer versions 7 and 8 (see Haddad et al., 2022), but there is overall more consistency between 7 and 8 than between prior version changes. This reinforces the need for caution when pooling data across studies processed using different versions of FreeSurfer.

Finally, automated processing pipelines, while invaluable, should not be treated as infallible. Researchers are encouraged to carefully review pipeline documentation, adhere to usage guidelines, and implement checks. It is crucial to remain methodologically vigilant when analyzing neuroimaging data. Although FreeSurfer 8 substantially improves automated template registration, this version is just now entering preprocessing workflows, and has not yet been integrated into fMRIPrep at the time of this writing.

### Recommendations

Aside from visual inspection of the registration step, there are several approaches researchers should consider to address the reliability concerns with eTIV and different FreeSurfer versions. First, when raw imaging data are available, researchers should process or re-process the data with Freesurfer 8 or later versions, particularly when eTIV is a key outcome or covariate, but also given continued reliability concerns. For longitudinal datasets, visually or quantitatively assessing change between scan sessions can provide an indication of measurement instability within an individual. With as few as two time points, percent change in eTIV can be used to identify potentially implausible changes relative to the population and interval being studied; in our data, we considered mean absolute changes exceeding 2% to indicate potential instability, as this exceeded the longitudinal stability observed with AssemblyNet. In addition to, or if longitudinal data is not available, calculating a whole brain volume to intracranial volume ratio may also help identify implausible results. Researchers may further consider comparing eTIV with alternate estimates of intracranial volume derived from segmentation based intracranial volume estimates such as sbTIV from SAMSEG in FreeSurfer versions prior to 8. Although segmentation-based estimates are known to generate systematic bias, such as producing larger values than scaling factor estimates, the correlation between sbTIV and eTIV should still be very strong (Nerland et al., 2022). Finally, researchers should examine the distribution of eTIV within their sample, including extreme values and unexpected clustering by age, sex, site, scanner, or processing version. Such patterns may indicate systematic processing differences or reconstruction failures that would not necessarily be apparent from individual quality metrics alone.

Moving forward, researchers should carefully evaluate ICV estimates, consider alternative brain size correction strategies, and conduct sensitivity analyses to ensure that developmental inferences are not driven by unstable global brain measures. Additionally, as these errors may have materially contaminated prior reports of developmental trajectories relying on automated FreeSurfer outputs, the field should judiciously reconsider our previous understandings and approach new findings derived from more reliable methods with an open mind.

## Acknowledgments

JHP, ARMS, AJR, and JS received support from NIH grants R01/R56 MH107418 and R01 MH127408, which enabled data acquisition and processing for this publication. ARMS and MNG received support from the National Center for Advancing Translational Sciences of the National Institutes of Health under award number TL1TR002371, which enabled data processing for this publication. AJR received support from NIH grant F32 MH144791. BTN and JVH received support from NIH grant S10 OD038249 which enabled data processing for this publication. The content is solely the responsibility of the authors and does not necessarily represent the official views of the National Institutes of Health.

## Supplementary Data

**Figure S1.**
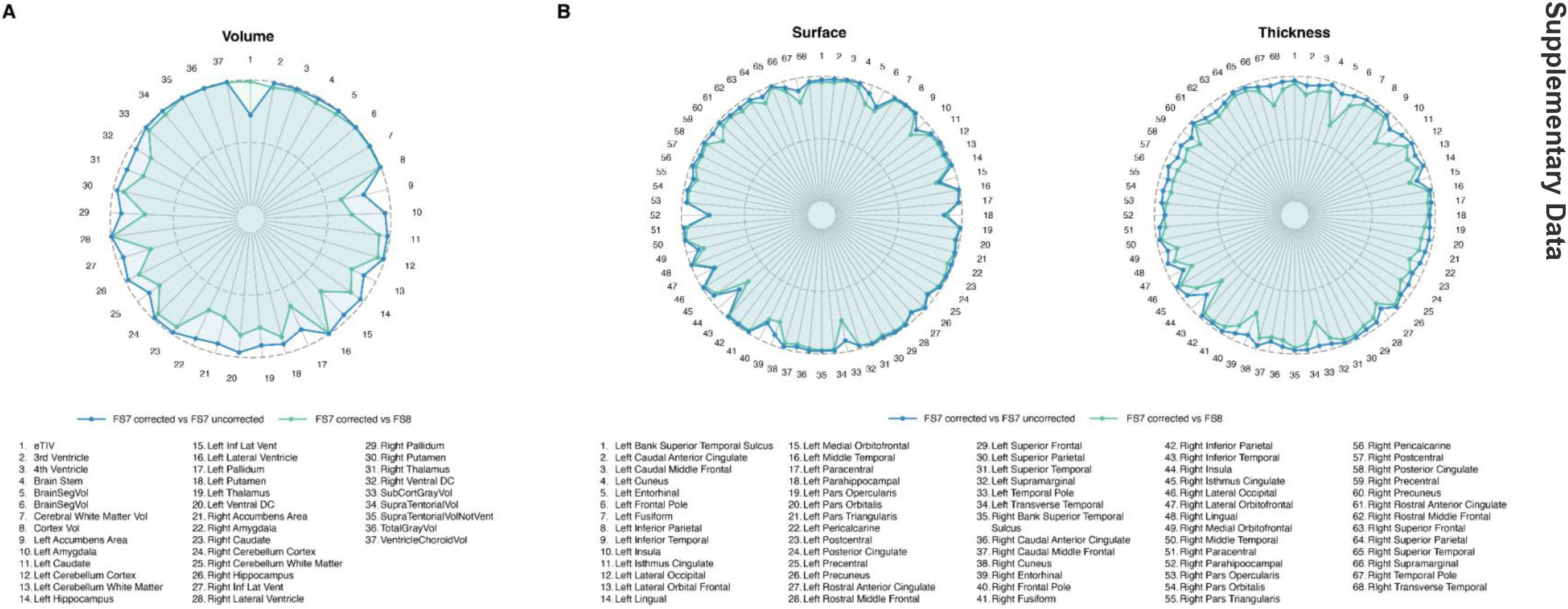
Regional interversion agreement (Pearson’s r) for corrected and uncorrected registration in FreeSurfer 7 (blue) and corrected FreeSurfer 7 and FreeSurfer 8 (green) for (A) volume and (B) surface and thickness. Concentric circles represent Pearson’s r at 0.0, 0.5, and 1.0.

