## Supplemental Figure 1 for "Widely Used Calculations of Intracranial Volume Provide Inaccurate Estimates in Developmental Cohorts: Evidence and Recommendations for the Neuroimaging Community"

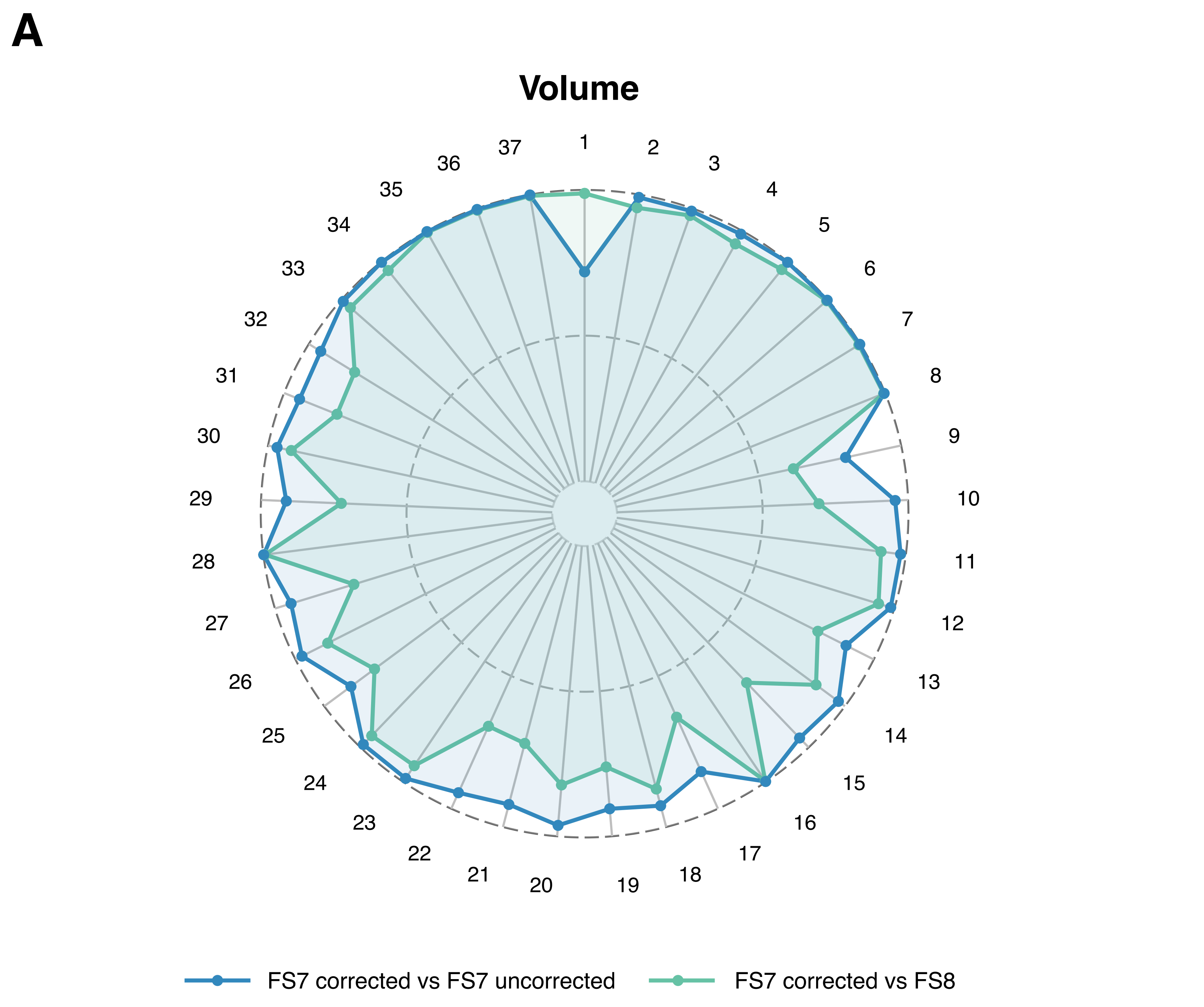


1. eTIV
2. 3rd Ventricle
3. 4th Ventricle
4. Brain Stem
5. BrainSegVol
6. BrainSegVol
7. Cerebral White Matter Vol
8. Cortex Vol
9. Left Accumbens Area
10. Left Amygdala
11. Left Caudate
12. Left Cerebellum Cortex
13. Left Cerebellum White Matter
14. Left Hippocampus
15. Left Inf Lat Vent
16. Left Lateral Ventricle
17. Left Pallidum
18. Left Putamen
19. Left Thalamus
20. Left Ventral DC
21. Right Accumbens Area
22. Right Amygdala
23. Right Caudate
24. Right Cerebellum Cortex
25. Right Cerebellum White Matter
26. Right Hippocampus
27. Right Inf Lat Vent
28. Right Lateral Ventricle
29. Right Pallidum
30. Right Putamen
31. Right Thalamus
32. Right Ventral DC
33. SubCortGrayVol
34. SupraTentorialVol
35. SupraTentorialVolNotVent
36. TotalGrayVol
37. VentricleChoroidVol

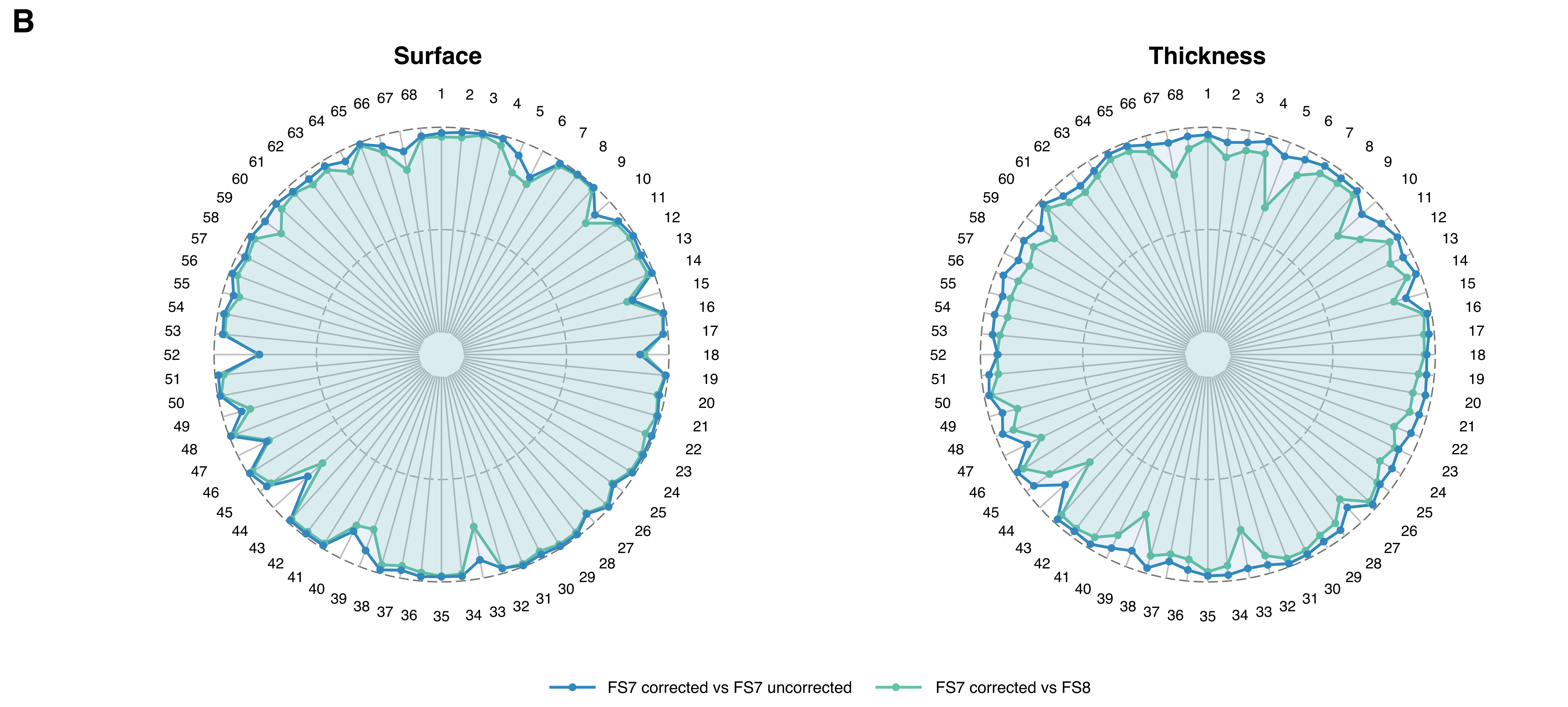


1. Left Bank Superior Temporal Sulcus
2. Left Caudal Anterior Cingulate
3. Left Caudal Middle Frontal
4. Left Cuneus
5. Left Entorhinal
6. Left Frontal Pole
7. Left Fusiform
8. Left Inferior Parietal
9. Left Inferior Temporal
10. Left Insula
11. Left Isthmus Cingulate
12. Left Lateral Occipital
13. Left Lateral Orbital Frontal
14. Left Lingual
15. Left Medial Orbitofrontal
16. Left Middle Temporal
17. Left Paracentral
18. Left Parahippocampal
19. Left Pars Opercularis
20. Left Pars Orbitalis
21. Left Pars Triangularis
22. Left Pericalcarine
23. Left Postcentral
24. Left Posterior Cingulate
25. Left Precentral
26. Left Precuneus
27. Left Rostral Anterior Cingulate
28. Left Rostral Middle Frontal
29. Left Superior Frontal
30. Left Superior Parietal
31. Left Superior Temporal
32. Left Supramarginal
33. Left Temporal Pole
34. Left Transverse Temporal
35. Right Bank Superior Temporal Sulcus
36. Right Caudal Anterior Cingulate
37. Right Caudal Middle Frontal
38. Right Cuneus
39. Right Entorhinal
40. Right Frontal Pole
41. Right Fusiform
42. Right Inferior Parietal
43. Right Inferior Temporal
44. Right Insula
45. Right Isthmus Cingulate
46. Right Lateral Occipital
47. Right Lateral Orbitofrontal
48. Right Lingual
49. Right Medial Orbitofrontal
50. Right Middle Temporal
51. Right Paracentral
52. Right Parahippocampal
53. Right Pars Opercularis
54. Right Pars Orbitalis
55. Right Pars Triangularis
56. Right Pericalcarine
57. Right Postcentral
58. Right Posterior Cingulate
59. Right Precentral
60. Right Precuneus
61. Right Rostral Anterior Cingulate
62. Right Rostral Middle Frontal
63. RIght Superior Frontal
64. Right Superior Parietal
65. Right Superior Temporal
66. Right Supramarginal
67. Right Temporal Pole
68. RIght Transverse Temporal
